# Expanding the Promoter Toolbox for Metabolic Engineering in the Lignocellulolytic Thermophile *Anaerocellum bescii*

**DOI:** 10.64898/2026.06.21.733613

**Authors:** Joey L. Galindo, Hansen Tjo, Archit Srivastava, Miranda Harmon-Smith, Ian Blaby, Jonathan M. Conway

## Abstract

*Anaerocellum* (formerly *Caldicellulosiruptor*) *bescii,* an anaerobic, extremely thermophilic (T_opt_ ∼78 °C) lignocellulolytic bacterium, is a promising chassis for metabolic engineering and next-generation bioprocessing. Yet, a lack of well-characterized genetic parts in *A. bescii* has hampered metabolic engineering efforts. Here, using a previously developed hyperthermophilic β-galactosidase reporter system, we screened a diverse panel of putative *A. bescii* promoter sequences, identifying promoters that drove reporter output across a broad range. For a select subset, we mapped their transcriptional start sites (TSSs) and evaluated ribosome binding site (RBS) regions using chimeric promoter constructs. By constructing truncated promoter variants, we defined functional regions within the widely used, high-expression S-layer protein promoter (P_slp_) and engineered a compact 99 bp variant that retained substantial reporter activity. Finally, we demonstrated that these new promoters can be used for metabolic engineering by using two newly characterized promoters to express an established thermostable alcohol dehydrogenase from *Thermoclostridium stercorarium* to drive ethanol production in *A. bescii*. Together, this work expands and diversifies the *A. bescii* genetic toolkit, opening doors to future metabolic engineering efforts in this species.

## Introduction

Non-model microorganisms often possess unique advantages over conventional biotechnology hosts, such as the ability to grow in extreme environments or catabolize recalcitrant feedstocks like lignocellulosic plant biomass. These traits make them promising new platforms for metabolic engineering^1–3^. However, harnessing this potential via synthetic biology requires a set of well-characterized robust genetic parts (e.g., promoters, ribosome binding sites, terminators), which are typically lacking in contrast to model organisms like *Saccharomyces cerevisiae* or *Escherichia coli*^1,2,4^. One such non-model species of interest for bioprocessing applications, *Anaerocellum* (formerly *Caldicellulosiruptor*) *bescii*, is amongst the most lignocellulolytic and thermophilic bacteria known, with an optimal growth temperature of 75-78 ℃ under anaerobic conditions^3,5,6^. Previous work developed a set of tools for genetic manipulations in *A. bescii*, including a highly thermostable kanamycin resistance gene (*htk*) for kanamycin selection and Δ*pyrF* or Δ*pyrE* uracil auxotroph strains for positive selection of uracil prototrophy and counterselection on 5-FOA^7,8^. These genetic tools have enabled metabolic engineering of *A. bescii* to produce several industrially relevant products including acetone, 2,3-butanediol, and ethanol^9–13^.

Developing a diverse library of characterized promoters is an important next step to expanding the genetic toolkit in *A. bescii*^6,14^. In non-model organisms, native promoters are often chosen as initial candidates for characterization, an approach taken in other thermophilic anaerobic bacteria like the more moderately thermophilic lignocellulolytic *Acetivibrio thermocellus* (formerly *Clostridium thermocellum*) or the acetogen *Thermoanaerobacter kivui*^2,15–19^. Yet, the toolbox of promoters in *A. bescii* remains especially limited, posing a major engineering bottleneck^5,6,14^. To date only four native promoters have been used for genetic engineering in *A. bescii*^7,9–13,20^. These include three constitutive promoters for the S-layer protein (P_slp_; Athe_2303), an S30 ribosomal protein (P_S30_; Athe_2105), and a bifurcating hydrogenase (P_bh_; Athe_1295), all of which are thought to drive strong constitutive expression. And, while they support functional expression *in vivo*, they have not been systematically benchmarked. A single inducible promoter has also been identified associated with the xylose isomerase gene (P_xi_; Athe_0603) that drives increased expression in response to the presence of xylose^20^. Moreover, metabolic engineering in *A. bescii* to date has almost exclusively relied upon P_slp_ to drive heterologous enzyme expression^6,9–13^.

Rapid characterization of genetic parts is enabled by readily observable and quantifiable protein-based reporter systems^2,5,21,22^. However, most conventional reporters are not compatible with the combination of extremely thermophilic and anaerobic growth conditions required by *A. bescii*. To address these issues in *A. bescii*, we previously developed a novel reporter system based on heterologous expression of a hyperthermophilic β-galactosidase (*Cm*βgal) from the archaeon *Caldivirga maquilingensis* ^22,23^. Using this *Cm*βgal-based reporter system, we evaluated a diverse panel of 19 native *A. bescii* promoters and benchmarked their strength against three known promoter sequences. To evaluate the effect of carbon source and growth phase, we tested the reporter expression driven by a subset of these promoters under various growth conditions. Additionally, we determined the putative transcriptional start sites (TSSs) for six promoter sequences via 5’ rapid amplification of cDNA ends (5’ RACE). This allowed us to map the transcriptional topology of P_slp_ in detail and characterize the ribosome binding sites (RBS) of several other promoters by expressing *Cm*βgal with truncated or chimeric promoter sequences. Finally, as a proof of concept for their utility for metabolic engineering, we demonstrated that two of these new promoters can be used to express a previously utilized AdhE from the bacterium *Thermoclostridium stercorarium* (*Tster*) and enable ethanol production in *A. bescii*. Together, this work expands the genetic toolset available in *A. bescii* by defining native and engineered promoters with a range of expression strengths and demonstrating their ability to enable heterologous product formation. By broadening the expression levels available for heterologous gene expression, these parts advance *A. bescii* as a platform organism for metabolic engineering and bioprocessing applications.

## Results

### Identification and Characterization of Native *A. bescii* Promoters

To develop a versatile suite of promoters for use in *A. bescii*, 19 native promoters were identified for testing alongside the three established promoters used in prior *A. bescii* genetic engineering (**Table S1**). These putative promoter sequences were defined as the 200 base pairs immediately upstream of the start codon in an associated gene in the *A. bescii* genome^24^. Promoter sequences were chosen from genes with varied transcriptional levels based on published transcriptomic data with the goal of capturing a broad range of potential levels of protein expression^25,26^. This selection of new promoters was also biased toward genes of known function, such as glycoside hydrolase enzymes and sugar transporters^25–28^. The promoter sequences were inserted ahead of the *Cm*βgal reporter^22^ on a replicating vector (**Fig. 1a**) and transformed into wild type *A. bescii.* Two control plasmids were also transformed, an empty vector control that contains no reporter gene, and a no promoter control which contains the *Cm*βgal gene but lacks an upstream promoter sequence entirely (**Fig. 1a**). In total 24 strains encompassing the two controls, 19 new promoters, and 3 characterized promoters (P_slp_, P_bh_, and P_xi_) were constructed.

**Figure 1.**
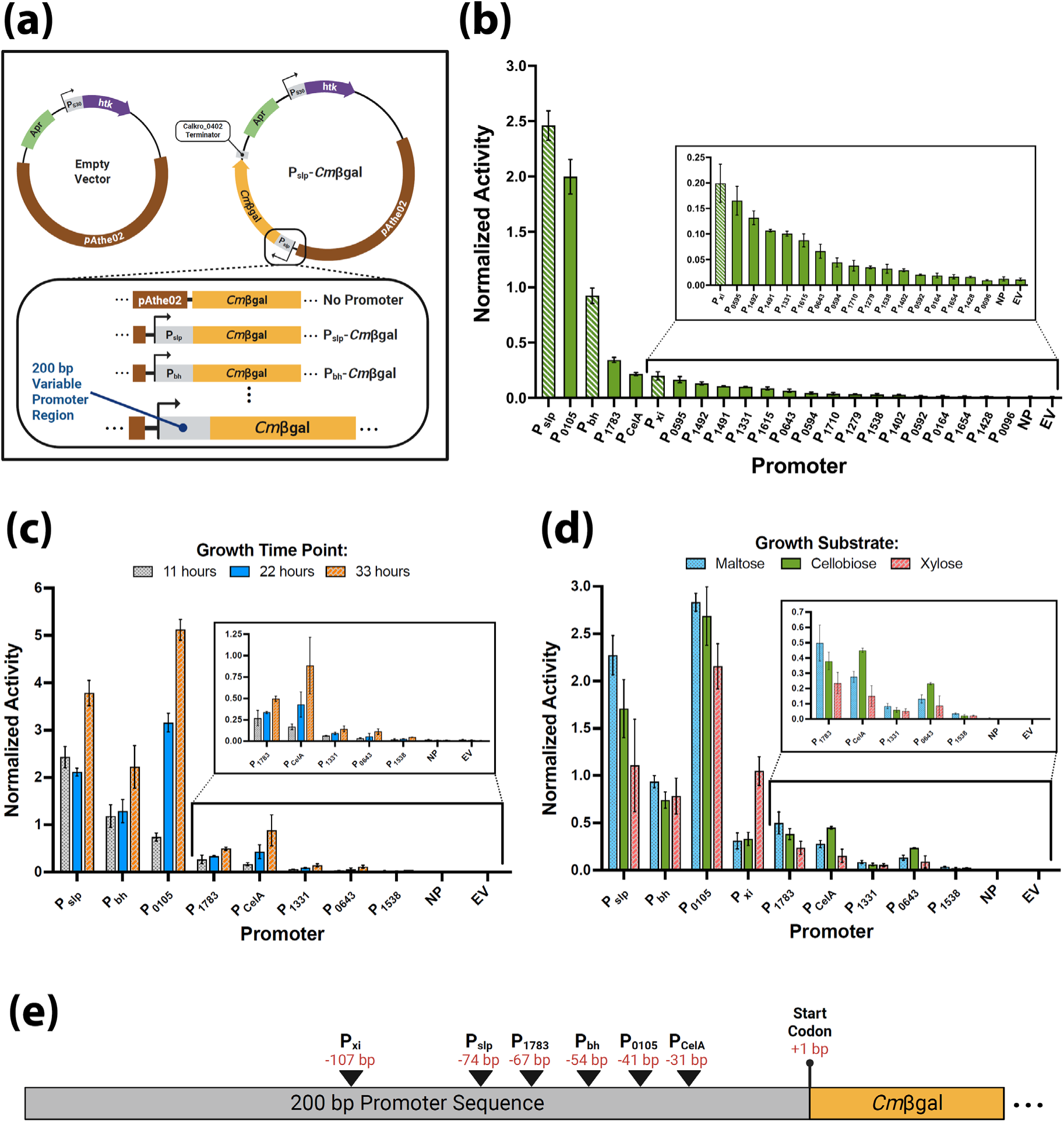
Characterization of native *A. bescii* promoters. **(a)** Plasmid maps of the *Cm*βgal reporter constructs transformed into *A. bescii* including as specific examples: P_slp_ (pJLG093), P_bh_ (pJLG161), the no promoter control (pJLG159), and the empty vector control (pSBS4). A complete list of plasmids and the promoters they contain is available in supplementary **Table S1** (**b**) Normalized β-galactosidase activity of the *A. bescii* strains containing various 200bp promoters expressing the *Cm*βgal reporter (Table S1). The 3 previously characterized promoters (P_slp_, P_bh_, and P_xi_) are highlighted with white stripes. **(c)** Normalized activity of a selection of 8 *Cm*βgal expressing *A. bescii* strains over the course of 33 hours of growth, with measurements taken at 11, 22, and 33-hour time points. **(d)** Normalized activity of the *A. bescii* strains tested in **(c)** as well as the P_xi_–*Cm*βgal strain grown on 3 types of defined media containing maltose, cellobiose, or xylose as the growth substrate. Error bars in panels (**b-d**) represent one standard deviation among biological triplicates. Supplementary data associated with panels (**b-d**) can be found in **Tables S2-4**. **(e)** Diagram indicating the location of transcriptional start sites within the 200bp promoter sequence determined via Sanger sequencing of 5’ RACE products (**Supplementary File S1**) for 6 promoters. Panels **a** & **e** were created in BioRender: Galindo, J. (2026) https://BioRender.com/21ng433 & Galindo, J. (2026) https://BioRender.com/lv4ojt9.

These strains were grown in biological triplicate on complex media with maltose as a carbon source and kanamycin selection (CM516K media) and the β-galactosidase activity was evaluated (**Fig 1b & Table S2**). Normalized β-galactosidase activity as described previously^22^, is defined as the net absorbance at 405 nm produced from the colorimetric para-nitrophenol-β-D-galactopyranoside (pNPβGal) substrate divided by the density (OD_680_) of the prepared cell input. In this assay, the benchmark P_slp_ and P_bh_ promoters showed the expected relative expression, with P_bh_ showing ∼40% the activity of P_slp_, as has been observed previously^9,22^. Most candidate promoters tested here drove lower expression of less than 14% that of the P_slp_ benchmark (**Fig. 1b**). The notable exception was P_0105_, which showed expression almost as high as P_slp_ (**Fig. 1b**). This promoter is associated with the gene for the substrate binding domain of an ABC sugar transporter (**Table S1**)^25,26^. This screen also revealed that P_1491_ and P_1492_ produce similar levels of expression from a 151 bp intergenic region between the divergently oriented genes, suggesting this functions as a bi-directional promoter (**Table S1** & **Fig. 1b**).

Next, to explore how growth stage and carbon source affect promoter activity, we selected a subset of promoters spanning a range of expression levels observed in the initial screen. This panel included the well-established benchmarks P_slp_ and P_bh_, the strong promoter P_0105_, moderate promoters P_1783_ and P_CelA_, and weaker promoters P_1331_, P_0643,_ and P_1538_ (**Table S1**). P_CelA_ was included because it is associated with the large-multidomain cellulase CelA, the most well studied *A. bescii* glycoside hydrolase enzyme^6,28,29^.

To see how promoter strength varied over the course of growth, *A. bescii* strains with each of these eight promoter–*Cm*βgal plasmids as well as the no promoter and empty vector control plasmids were grown in triplicate for 33 hours on CM516K media. Cultures were sampled after 11, 22, and 33 hours of growth, roughly corresponding to mid-exponential, late-exponential, and early stationary phase growth, respectively (**Fig. 1c & Table S3**). Observed activity from all the tested promoters increased as cells entered stationary phase after 33 hours of growth, while the relative expression levels among the promoters were generally preserved (**Fig. 1c & Table S3**). However, the most dramatic increase was observed for P_0105_, with its activity increasing from approximately 30% to 140% that of P_slp_ between the 11 and 33-hour time points (**Fig. 1c**).

Next, to see if promoter output was affected by carbon source, these same strains, along with P_xi_ as a known xylose-inducible promoter, were grown in triplicate on defined media containing one of three common *A. bescii* substrates: maltose (DM516K media), cellobiose (DC516K media), or xylose (DX516K media) (**Fig. 1d & Table S4**). Generally, activity was highest on maltose relative to cellobiose or xylose, with P_slp_ showing a particularly notable decrease on the latter two substrates (**Fig. 1d**). In contrast, P_CelA_ and P_0643_ showed increased activity on cellobiose (**Fig. 1d**). As expected, reporter activity from P_xi_ was higher on xylose than on cellobiose or maltose, increasing 3.15x and 3.35x, respectively, while remaining high on the non-xylose substrates compared with most other promoters (**Fig. 1d**). While reporter output is normalized to the OD680 input to the assay, strains generally reached higher densities on xylose and cellobiose (OD680 ∼0.21) than maltose (OD680 ∼0.09) in these assay conditions **(Table S4)**. Across these assays (**Fig 1b, c, d**), both the empty vector and no promoter control strains showed no β-galactosidase activity. For subsequent reporter assays, the no promoter strain was used as the preferred negative control.

Finally, to explore where transcription initiates within these 200 bp promoter sequences, RNA was extracted from strains expressing *Cm*βgal from P_slp_, P_bh_, P_0105,_ P_1783_, P_CelA_, and P_xi_. Utilizing a template switching oligonucleotide (TSO)-based 5’ rapid amplification of cDNA ends (5’ RACE) approach, we determined the putative transcriptional start sites (TSSs) of these promoter sequences (**Fig. 1e & Supplementary File S1**). Alignment of Sanger-sequenced 5’ RACE products with their associated reporter vector revealed TSSs ranging from 31 bp upstream of the start codon for P_CelA_ to 107 bp upstream of the start codon for P_xi_ (**Fig. 1e**).

### Mapping functional elements within the P_slp_ promoter

The P_slp_ promoter has historically been the primary promoter used for metabolic engineering in *A. bescii* ^6,9–13^. Given its widespread use, we used the mapped TSS at the -74 bp position (**Fig. 1e**) to guide a truncation-based functional analysis of this promoter. Truncations targeting the predicted transcriptional elements (-35 and -10 box) and 5’ untranslated region were constructed upstream of the *Cm*βgal reporter and transformed into *A. bescii* (**Fig. 2a** & **Table S1**). The 5’ truncations P_slp127,_ P_slp108_, and P_slp89_ progressively removed sequence ahead of the TSS, with P_slp108_ and P_slp89_ truncating into the -35 and -10 transcriptional elements, respectively. P_slp173_ removed 27 bp between the TSS and the predicted Shine-Dalgarno sequence^30,31^ (**Fig. 2a**). P_slp100_ was designed as a minimal functional P_slp_ sequence by combining the truncations of P_slp127_ and P_slp173_ to generate a shorter promoter variant retaining the predicted core transcriptional (-35 and - 10 boxes) and translational (Shine-Dalgarno) elements (**Fig. 2a**). Sequence verification of the P_slp100_ construct from the final *A. bescii* strain revealed a single “a” base deletion at the -28 bp position, therefore this promoter was designated P_slp99_.

**Figure 2.**
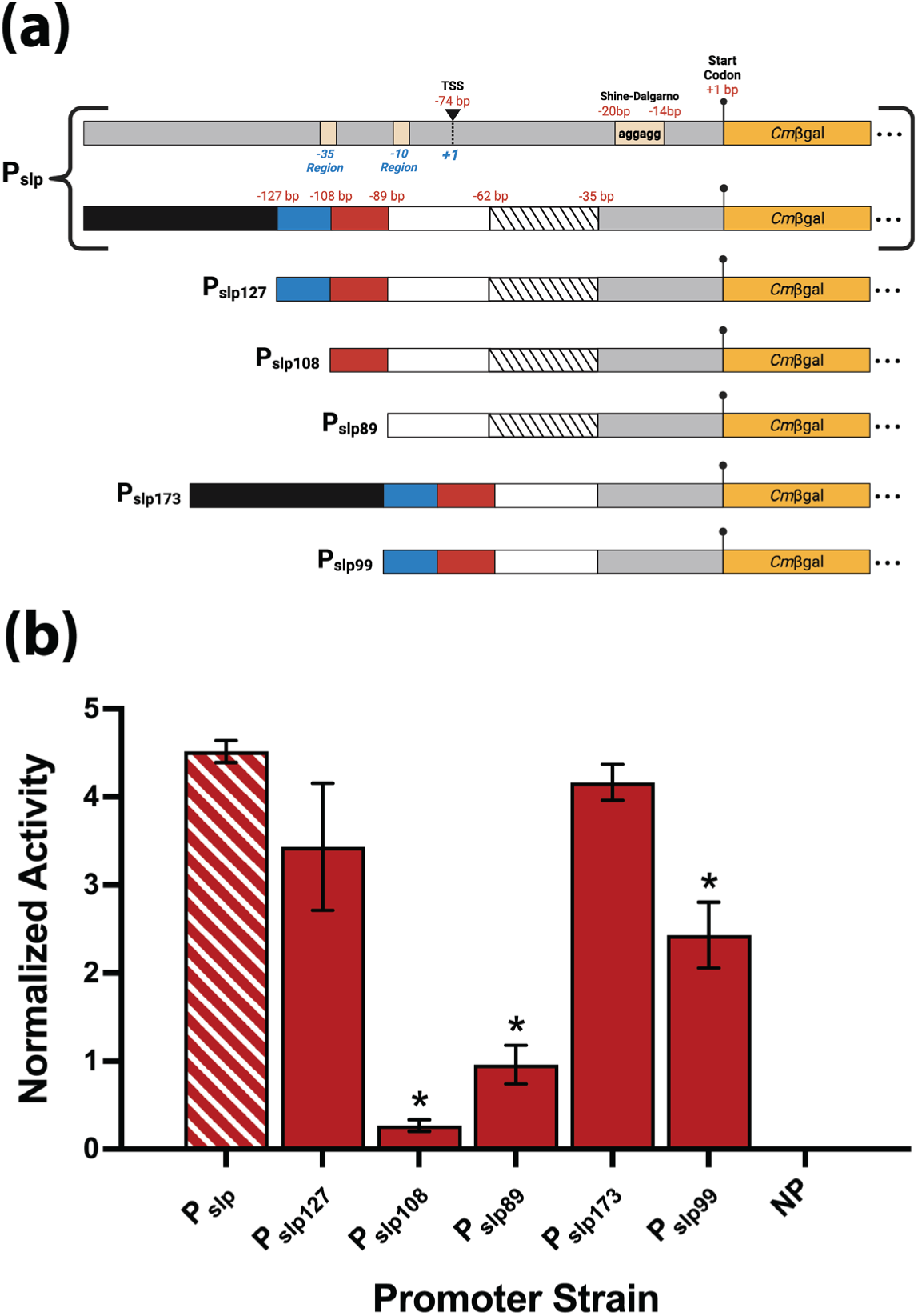
Constructing and testing truncations of the S-layer protein promoter (P_slp_). **(a)** Diagram outlining the major features of P_slp_ and the truncated variants that were created (**Table S1**; pJLG215-19). Red numbers indicate locations relative to the start codon while blue numbers in italics indicate locations relative to the TSS. **(b)** Normalized β-galactosidase activity of prepared *A. bescii* cells grown on CM516K medium expressing the *Cm*βgal reporter protein with the truncated P_slp_ variants described in **(a)** alongside the no promoter (NP) control. White stripes indicate the strain with the native P_slp_ promoter. Asterisks indicate statistically significant (*P* ≤ 0.05) differences in activity compared with full-length P_slp_ construct determined via a Brown-Forsythe and Welch ANOVA test, with resulting *P*-values of 0.3435, <0.0001, 0.0006, 0.2862, and 0.0354 for comparison to P_slp127_, P_slp108,_ P_slp89_, P_slp173_, and P_slp99_, respectively. Error bars represent one standard deviation between biological triplicates. Supplementary data associated with panel **b** can be found in **Table S5**. Panel **a** was created in BioRender. Galindo, J. (2026) https://BioRender.com/ dtyoo3n.

The *A. bescii* strains containing these P_slp_ truncation-*Cm*βgal constructs were grown in triplicate on CM516K media alongside the full-length P_slp_ and no promoter control, and reporter output was evaluated (**Fig. 2b & Table S5**). P_slp127_ and P_slp173_ drove reporter activity that was not statistically different (*P* > 0.05) from the full-length P_slp_ sequence, indicating that removal of either the distal upstream region or 27 bp region between the TSS and predicted RBS did not significantly reduce expression. In contrast, P_slp108_ and P_slp89_ drove significantly lower expression than full-length P_slp_, reaching only 6% and 21% of P_slp_ activity, respectively, consistent with disruption of the core -35 and -10 promoter elements. The combined truncation in P_slp99_ created a compact 99 bp promoter that retained 54% the reporter activity of full-length P_slp_, demonstrating that a compact Pslp-derived promoter retaining the predicted core promoter elements can still drive substantial expression (**Fig. 2b**).

To determine whether the truncations altered transcription initiation, RNA was extracted from the same *A. bescii* strains and analyzed by 5’ RACE (**Supplementary File S1**). Sequencing of these products indicated that transcription from P_slp127_, P_slp173_, and P_slp99_ is initiated from the same sequence position as identified in the full-length Pslp. Conversely, for the weak expression truncations, 5’ RACE on P_slp108_ produced a mixture of possible TSSs, and P_slp89_ had a putative TSS 188 bp upstream of the promoter sequence within the native *pAthe02* plasmid backbone^7,8^. This upstream TSS may explain the statistically significant (*P*= 0.024) increase in β-galactosidase activity observed for P_slp89_ relative to P_slp108_ (**Fig. 2b**). Interestingly, although the 5’ RACE sequencing quality was poor for the no promoter control, the sequencing suggested low-level transcription from a similar region of the plasmid backbone as P_slp89_ despite the absence of detectable *Cm*βgal activity in the no promoter control.

### Characterization of ribosome binding sites from native *A. bescii* promoters

Because both transcriptional and translational elements contribute to overall protein expression, next we examined putative ribosome binding site (RBS) regions from select native promoters in a common transcriptional context. RBS sequences are commonly used to coordinate translation from polycistronic operons in bacteria, and putative RBS sequences have previously been used for operon design in *A. bescii* despite not being individually characterized ^9,11^. We therefore tested RBS regions from native promoters whose TSSs had been mapped by 5’ RACE. For this analysis, we defined the putative RBS region as the 30bp immediately upstream of the start codon (**Fig. 3a**). To test these RBSs while holding the upstream sequence driving transcription constant, we fused the 5’ 170bp of P_slp_ to the 30 bp putative RBS regions from P_bh_, P_0105,_ P_1783_, and P_CelA_, generating chimeric promoters termed P_slp_::RBS_bh_, P_slp_::RBS_0105_, P_slp_::RBS_1783_, and P_slp_::RBS_CelA_ respectively (**Table S1** & **Fig. 3a**).

**Figure 3.**
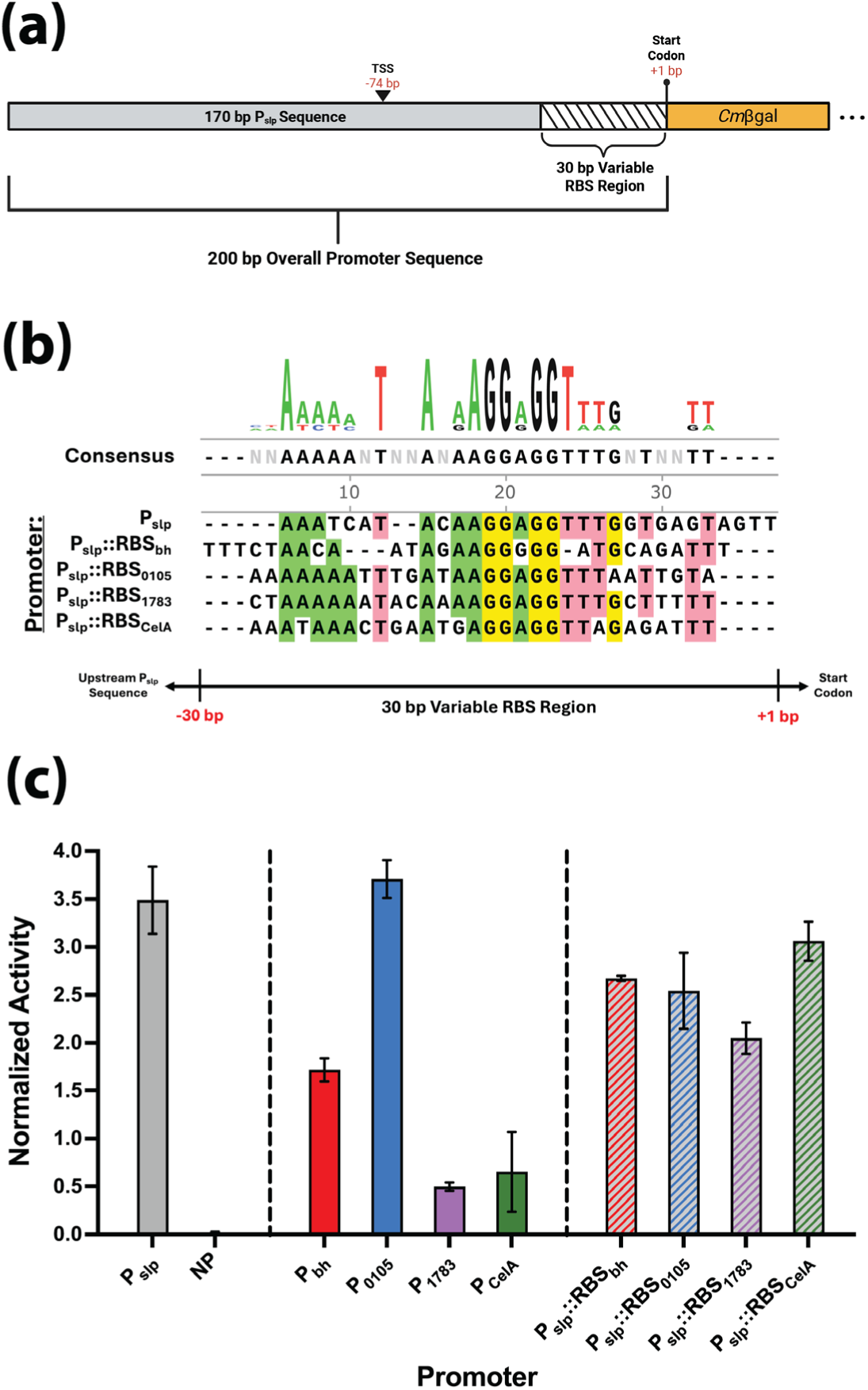
Using chimeric promoter sequences to characterize ribosome binding sites (RBS) in *A. bescii*. **(a)** Diagram of the reporter constructs (**Table S1;** pJLG231-34) expressing the *Cm*βgal reporter gene with 200 bp chimeric P_slp_::RBS sequences that consist of the 5’ 170bp of P_slp_ combined with the 30bp immediately upstream of the start codon corresponding to the putative RBS of 4 other native promoters: P_bh_, P_0105_, P_1783_, and P_CelA_. **(b)** Multiple sequence alignment of the four chimeric promoters with P_slp_ using ClustalOmega^32^. The upstream sequence of all five promoters is the identical native P_slp_ sequence while variability can be seen in the 30bp RBS region shown, though all 5 RBSs contain a consensus (AGGAGG or AGGGGG) Shine-Dalgarno sequence 10-14bp ahead of the start codon. **(c)** Normalized β-galactosidase activity of prepared *A. bescii* cells grown on CM516K medium expressing *Cm*βgal with the four chimeric promoters, the five original native promoters from which these chimeric promoters were constructed, as well as a no promoter (NP) control. Striped bars indicate chimeric promoter sequences. Error bars represent one standard deviation between biological triplicates. Supplementary data associated with panel **c** can be found in **Table S6**. Panel **a** was created in BioRender. Galindo, J. (2026) https://BioRender.com/xfp7662.

Multiple sequence alignment of these putative RBS regions using ClustalOmega^32^ showed that while there is considerable sequence variability across the 30bp, most contained a consensus AGGAGG Shine-Dalgarno motif, with P_bh_ containing an AGGGGG variant (**Fig. 3b**). The spacing between this motif and the ATG start codon was 10bp for all of the RBSs, except P_slp_ where this space was 14bp (**Fig 3b**). *A. bescii* strains with the chimeric Pslp::RBS-*Cm*βgal constructs (**Table S1** & **Fig. 3a**), the no promoter control, as well as the corresponding native promoter constructs were grown in biological triplicate in CM516K medium and evaluated using the β-galactosidase reporter assay (**Fig. 3c & Table S6**). The native promoter constructs showed reporter outputs similar to what was observed above (**Fig. 1b-d**), whereas the chimeric constructs produced outputs more similar to P_slp_ rather than their RBS source promoter (**Fig. 3c**). For example, native P_1783_ and P_CelA_ produced 14% and 19% of the activity of P_slp_, while P_slp_::RBS_1783_ and P_slp_::RBS_CelA_ show much higher activity at around 59% and 88% that of P_slp_ (**Fig. 3c**). These results suggest that the variation in observed protein expression between these five native promoters is due primarily to differences in transcription rather than translation. Furthermore, all of these 30bp RBS sequences appear to enable protein translation, indicating they may be useful for deployment in other genetic contexts such as polycistronic operon construction.

### Utilizing new promoters to drive ethanol production in *A. bescii*

Finally, to test whether these newly characterized promoters could drive product formation in *A. bescii*, we selected P_0105_ and P_1331_ to express the alcohol dehydrogenase (AdhE) from *Thermoclostridium stercorarium* (*Tster*). We used the *Tster* AdhE D492G variant, which has been used previously to maximize ethanol production in *A. bescii* and can utilize both NADH and NADPH as cofactors^13^. Replicating vectors based on the *Cm*βgal reporter constructs were modified to express codon-optimized *Tster AdhE_D492G* from either P_0105_ or P_1331_ and transformed into wild type *A. bescii* (**Fig. 4a**). These strains, along with the empty vector control, were grown in triplicate on complex medium with cellobiose as a carbon source and kanamycin selection (CC516K) at 65 ℃ for 48 hours (**Table S7**), conditions selected to support *Tster* AdhE_D492G activity and ethanol production^13^.

**Figure 4.**
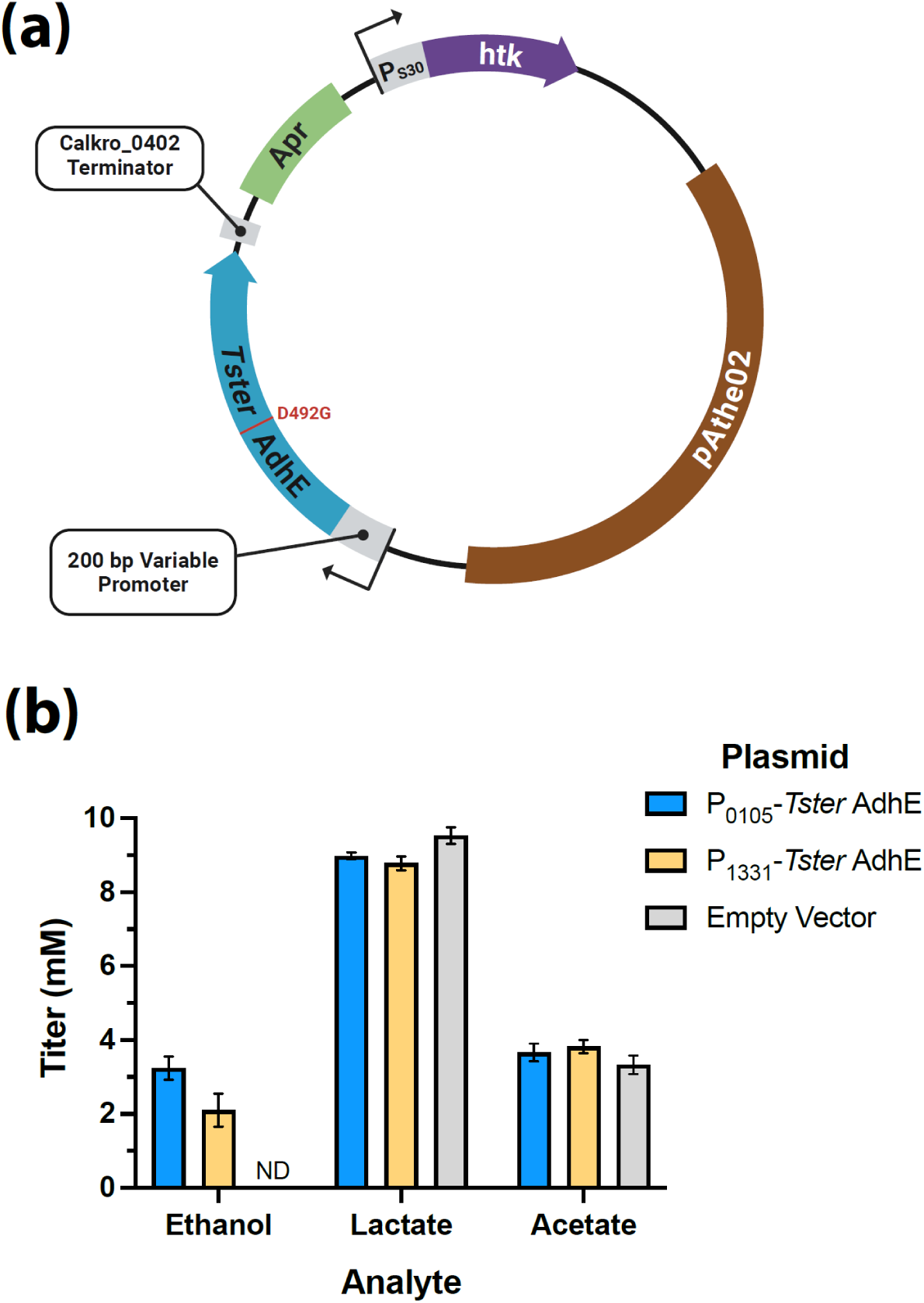
Ethanol production from *A. bescii* expressing the *Tster* AdhE with the D492G modification. **(a)** Map of the replicating vectors transformed into *A. bescii* used to express the *Tster* AdhE with P_0105_ or P_1331_ (**Table S1**; pJLG222 & 224). **(b)** Quantification of fermentation products from strains of *A. bescii* transformed with the *Tster* AdhE vectors described in **(a)** as well as the empty vector control after growth at 65 ℃ on CC516K medium. ND=not detected. Error bars represent one standard deviation among biological triplicates. Final OD680 cell densities of the *A. bescii* cultures associated with panel **b** can be found in **Table S7**. Panel **a** was created in BioRender. Galindo, J. (2026) https://BioRender.com/c5×3mpb.

After growth, ethanol, acetate, and lactate were quantified via HPLC. Both P_0105_ and P_1331_ drove expression resulting in detectable ethanol production, while ethanol was not detected in the empty-vector control (**Fig. 4b**). The P_0105_ construct, which showed higher reporter output than P_1331_ in the *Cm*βgal assay, produced 3.2 mM ethanol compared with 2.1 mM from the P_1331_ construct (**Fig. 4b**). These ethanol titers are lower than those reported previously for genomically integrated *adhE* genes in *A. bescii*^9,12,13^. All strains produced acetate and lactate at titers of 3.3-3.8 mM and 8.8-9.5 mM, respectively (**Fig. 4b**). Together, these results demonstrate that the newly characterized promoters can support heterologous enzyme expression sufficient to drive detectable product formation in *A. bescii*.

## Discussion

In this study, we deployed the previously developed *Cm*βgal reporter system^22^ to benchmark a suite of new promoters in *A. bescii*. These promoters had varied strengths ranging from barely detectable above background to greater than that of the widely used P_slp_ (**Fig. 1b**). We then characterized a subset of these promoters across growth stages and on three different carbon sources (**Fig. 1c & d**). Reporter expression appeared to increase for all tested promoters as cells entered stationary phase, an effect we previously noted for P_slp_ and P_bh_ during the original development of the *Cm*βgal reporter^22^, though here this behavior was more pronounced for P_0105_ (**Fig. 1c**). Reporter activity was moderately higher for most promoters on maltose, with a few exceptions, including P_CelA_, which showed increased reporter activity on cellobiose (**Fig. 1d**). This pattern may reflect cellobiose-dependent upregulation of the glucan degradation locus (GDL), which encodes CelA and the other primary cellulases of *A. bescii*^25,28^. Additionally, as expected, P_xi_ showed much higher expression on xylose (3.15-3.35x) compared with cellobiose or maltose, although it still remained relatively active on non-xylose carbon sources (**Fig. 1d**). This was a far smaller dynamic range than the 32-fold increase reported previously by *Williams-Rhaesa et al.* for P_xi_-driven lactate dehydrogenase specific activity measured *in vitro* from cell-free extracts of an *A. bescii* Δ*ldh* strain carrying a chromosomally inserted P_xi_-*ldh* construct^20^. The reduced apparent dynamic range of P_xi_ in the *Cm*βgal reporter system may reflect differences between reporter assays, genomic versus plasmid-based expression, or growth dynamics across experiments. Because the turnover rate of the *Cm*βgal protein in *A. bescii* has not been determined and our activity output is normalized to cell density, reporter concentration, and by extension activity, may be diluted by more rapidly dividing cells in some growth conditions.

To further characterize select promoters, we utilized 5’ RACE to determine putative transcriptional start sites (TSSs) for six of the tested promoters. These TSSs ranged from as close as 31bp upstream of the start codon for P_CelA_ to as far as 107 bp upstream of the start codon for P_xi_ (**Fig. 1e**). This latter result agreed well with the location of a xylose regulator binding site proposed by *Williams-Rhaesa et al.* for P_xi_ between 76 and 99 bp upstream of the start codon^20^. With these putative TSSs in hand, we then mapped the transcriptional and translational topology of P_slp_ by using truncations of the native sequence to express *Cm*βgal (**Fig. 2a**). This allowed us to shorten the P_slp_ sequence to as little as 99bp and still retain >50% of *Cm*βgal reporter expression driven by the native promoter (**Fig. 2b**). Next, we tested putative 30bp ribosome binding sites (RBS) from P_bh_, P_0105,_ P_1783_, and P_CelA_ in chimeric promoters with P_slp_-driven transcription via its 5’ 170bp sequence (**Fig. 3a**). Expression of *Cm*βgal with these chimeric sequences showed activity much more comparable to that of P_slp_ than to the promoter source of the RBS (**Fig. 3c**), indicating that in this case differences in expression across these native promoters were not primarily driven by differences in ribosome binding efficiency. These results also validated the RBS sequences, which would enable their use in the future for the design of multi-enzyme polycistronic operons in *A. bescii*, as has been done previously^9,11^.

Finally, to demonstrate that these new promoters can be used for metabolic engineering and product formation, we utilized two of these new promoters, P_0105_ and P_1331_, to express the *Tster* AdhE_D492G enzyme used by *Bing et al.* for ethanol production in *A. bescii*^13^. Expression of AdhE with these promoters from a replicating vector in wild type *A. bescii* resulted in detectable ethanol production in both strains (**Fig. 4b**). However, while these titers were substantially lower than those reported for previously optimized ethanol-producing strains^9,11–13,33^, they represent the first reported use of promoters other than P_slp_ to drive non-native product formation in *A. bescii*^9–13^. We also observed relatively high lactate/acetate ratios in these cultures (**Fig. 4b**). This may reflect the lack of agitation during growth as accumulation of fermentative H_2_ in the liquid medium can shift metabolic flux toward lactate and ethanol production to balance the redox environment^13,34^.

Previously, only a handful of *A. bescii* promoters had been characterized, limiting the expression levels available for heterologous pathway design. Reflecting this limited toolkit, essentially all heterologous pathways in engineered *A. bescii* strains reported to date have relied on P_slp_^6,9–13^. Taken as a whole, this work has expanded the *A. bescii* genetic toolkit by characterizing native, chimeric, and engineered promoters that drive a range of expression levels in this host of interest for bioprocessing applications. In addition, by mapping the TSSs and validating modular RBS elements, this work has broadened the set of modular parts available for expression construct design in this organism. This expanded catalog of genetic parts is important for optimizing heterologous pathways because, as has been observed in other metabolic engineering applications, pathway optimization often requires balancing enzyme expression levels rather than simply maximizing expression^35–37^, and because certain heterologous proteins can be toxic when overexpressed^2^. These characterized parts may also inform genetic tool development in related thermophilic anaerobes, particularly given the prior use of P_xi_ as an inducible promoter in *Acetivibrio thermocellus* (formerly *Clostridium thermocellum*)^16^. By expanding the parts available for heterologous gene expression, this work strengthens the foundation for engineering *A. bescii* as a thermophilic platform for metabolic engineering and bioproduction.

## Materials and Methods

### Bacterial strains and culture maintenance

Plasmids were cloned in chemically competent *Escherichia coli* 10beta (New England Biolabs) or TOP10 (Thermo Scientific) and maintained as previously described^22^. Unless otherwise stated, *A. bescii* strains were cultured in 50 mL of C516 medium in 125 ml serum bottles sealed with 20 mm butyl rubber stoppers at 70 ℃ without shaking following the recipe described by Lipscomb *et al.* with the carbon source (5 g/L) indicated by a second letter of the medium identifier (e.g. CM516 medium contains maltose substrate)^7^. The C516 medium was supplemented with 50 µg/ml kanamycin (IBI Scientific) as appropriate and is referred to as C516K medium. Sealed serum bottles containing sterile-filtered media were made anaerobic through gas sparging, with the headspace being replaced with 80% (v/v) N_2_ and 20% (v/v) CO_2_ gas. Cell density of *A. bescii* cultures was measured as the optical density at 680 nm (OD680) using a cuvette in a Nanodrop One C spectrophotometer (Thermo Scientific) blanked with 1x DSM 516 salt solution.

### Vector and *A. bescii* strain construction

**Table S1** describes all the plasmids transformed in *A. bescii* in this study. Oligonucleotide primers and synthesized DNA used to construct the new plasmids for this study can be found in supplementary **Table S8** and **Table S9**, respectively. pJLG093 expressing *Cm*βgal with P_slp_ was constructed as described previously from the pSBS4 (empty vector) that was obtained from the lab of Dr. Robert Kelly (North Carolina State University)^7^. This vector consists of a native *A. bescii* replicating plasmid (*pAthe02*), the highly thermostable kanamycin resistance (*htk*) gene expressed by promoter P_S30_ associated with the S30 ribosomal protein (Athe_2105), as well as elements that enable cloning in *E. coli* including an apramycin resistance marker^7,8^ (Fig. 1a). Additionally, all of the Gibson assembly steps described below were carried out using the NEBuilder HiFi DNA assembly kit (New England Biolabs) as per the manufacturer’s instructions.

pJLG161-211 were constructed from pJLG093 in partnership with the Department of Energy Joint Genome Institute (JGI) at Lawrence Berkeley National Lab (Berkeley, CA) as described below. pJLG161-211 are identical to pJLG093 except that expression of *Cm*βgal is driven instead by a different 200 bp promoter sequence (Fig. 1a**, Table S1**). To create these vectors, pJLG093 was first modified to create unique PmeI sites, linearized by PCR amplification (**Table S8**; Primers JGI.P1 & JGI.P2), and re-circularized via Gibson assembly together with an ultramer (**Table S8**; Ultramer JGI.UM1) purchased from Integrated DNA Technologies to create the vector pJLG093_PmeI. After validation of this modified vector, sequences corresponding to new promoters flanked by linkers designed for assembly into pJLG093_PmeI linearized by PmeI digest (**Table S9**), were purchased (Twist Biosciences) and assembled. Several promoter sequences (**Table S9**; highlighted in red) contained synthesis constraint violations preventing their synthesis as single fragments. Instead, these were partitioned in silico using the BOOST suite^38^ of software tools and each purchased as two ultramers designed with internal Gibson assembly linkers. Using the same replicating vector backbone as the *Cm*βgal constructs described above, another plasmid named pJLG104 was constructed that contained the gene for the alcohol dehydrogenase (AdhE) from *Thermoclostridium stercorarium* (*Tster*) modified (D492G) as previously described^13^. The codon optimized gene for *Tster* AdhE_D492G (**Table S9**) was purchased (Twist Biosciences) and assembled into the unmodified pJLG093 backbone linearized without the *Cm*βgal gene (**Table S8**; Primers JGI.P3 & JGI.P4). Assemblies were transformed into chemically competent *E. coli* Top10 and candidate colonies were picked, sequence verified on the Pacific Biosciences Revio platform (Pacific Biosciences), and analyzed using custom pipelines at the Joint Genome Institute.

The following vectors were cloned in chemically competent *E. coli* 10beta and confirmed via whole plasmid sequencing (Azenta Genewiz, Plasmid-EZ). pJLG100, which is also identical to pJLG093 except expression of *Cm*βgal is driven by P_xi_, was constructed via Gibson assembly of the linearized pJLG093 backbone (**Table S8**; Primers JLG177 & 204) and the 200bp P_xi_ sequence amplified (**Table S8**; Primers JLG188 & 203) with overlaps from WT *A. bescii* genomic DNA (**Fig. 1a**). pJLG159 contains the *Cmβgal* gene with no upstream promoter sequence and was constructed from pJLG093 linearized via PCR amplification (**Table S8**; Primers JLG177 & 239) to delete the 200bp P_slp_ sequence entirely and re-circularized via Gibson assembly (**Fig. 1a**). pJLG215-219 express *Cm*βgal with various truncations of P_slp_ (**Fig. 2a**). pJLG215-218 were constructed from pJLG093 via PCR linearization (**Table S8**; Primers JLG177 & 245-49) to exclude parts of P_slp_ and re-circularization via Gibson assembly. pJLG219 was subsequently constructed in a similar manner (**Table S8**; Primers JLG248 & 249) from pJLG215. pJLG231-34 express *Cm*βgal with a 200 bp chimeric promoter sequence consisting of the 30bp immediately upstream of the start codon corresponding to the putative RBS sequences of P_bh_, P_0105_, P_1783_, and P_CelA_, respectively, combined with the remaining 170bp of P_slp_ (**Fig. 3a**). pJLG231-34 were constructed by first linearizing pJLG093 (**Table S8**; Primers JLG178 & 278) to exclude 30bp of P_slp_. Separately the 30bp RBS and *Cm*βgal gene from pJLG161, pJLG174, pJLG167, and pJLG168 were amplified with appropriate overlaps (**Table S8**; Primers JLG182 & 279-82) and Gibson assembled with the truncated pJLG093 backbone to create pJLG231-34 respectively. Finally, pJLG222 & 224 were constructed by linearizing pJLG174 & 208 (**Table S8**; Primers JLG178, 254, & 258) without the *Cm*βgal gene via PCR respectively and Gibson assembling these products with the *Tster* AdhE_D492G gene amplified from pJLG104 with appropriate overlaps (**Table S8**; Primers JLG180, 255, & 259).

For transformation into *A. bescii*, plasmid DNA was extracted from *E. coli* and prepared as previously described^7,22,39^. The parent for all constructed strains was wild type *A. bescii* DSM 6725 obtained from the lab of Dr. Robert Kelly (North Carolina State University). Competent *A. bescii* were grown on CM516 media supplemented with amino acids or tryptone (0.5 g/L) to an optical density at 680 nm (OD680) of 0.04-0.12, and transformations were carried out as described previously^7,22^. In brief, cells were washed in 10% (w/v) sucrose and transformed with 1-2 µg of methylated plasmid using a Bio-Rad gene pulser at 1800-2200 V, 400-600 Ω, and 25 µF. After recovery, transformed cells were selected for kanamycin resistance via growth in CM516K media. Cultures that grew in selective media were then plated and grown in solid CM516K media in an anaerobic chamber at 70 ℃ under a >98% (v/v) N_2_ and 1-2% (v/v) H_2_ atmosphere. Single colonies were picked and screened via colony PCR (**Table S8**; primers JLG181 & 224), with the presence of the correct promoter-reporter sequences confirmed by long-read sequencing of the colony PCR products (Azenta Genewiz, PCR-EZ).

### Growth conditions for testing *Cm*βgal expressing *A. bescii* strains

In the following experiments, with the exception of the growth curve test where cells were cultured in 125 mL serum bottles as described above, strains of *A. bescii* transformed with the empty vector and *Cm*βgal constructs were cultured in 18×125 mm “Balch-type” anaerobic tubes filled with 10 ml of degassed media and sealed with 20 mm butyl rubber stoppers in an anaerobic chamber with headspace containing >98% (v/v) N_2_ and 1-2% (v/v) H_2_. To start the growth of *A. bescii* transformed with the empty vector and *Cm*βgal constructs described above, pre-heated CM516K media was inoculated to a target OD680 range of 0.002-0.005 using cultures of *A. bescii* grown in the same media. The exception was the experiment testing the effects of different sugar substrates on promoter strength, for which inoculum cultures were grown on defined selective media that excludes yeast extract with maltose as the growth substrate, referred to as DM516K media. To begin the experiment these cultures were first washed in sugar-free defined selective media (D516K) as was described previously by *Tjo et al*^27^. These washed cells were then used to inoculate three types of defined selective media that each had different growth substrate: maltose (DM516K), cellobiose (DC516K), or xylose (DX516K). In all experiments, ∼5ml samples of grown *A. bescii* cells were harvested and frozen at -80 ℃ for subsequent β-galactosidase assays as previously described^22^. The OD680 of the source cultures of the samples were also measured.

For the initial screening of all native promoters, *A. bescii* transformed with *Cm*βgal expressing plasmids as well as the control vectors were grown for 18 hours at 70 ℃. For the test of different media on promoter expression, selected *Cm*βgal expressing *A. bescii* strains alongside the two controls were grown for 30 hours at 70 ℃ in DM516K, DC516K, or DX516K media. For the experiment testing the effect of growth on promoter expression, *A. bescii* transformed with these same plasmids except pJLG100 were grown for a total of 33 hours at 70 ℃. After 11, 22, and 33 hours of growth, OD680 measurements were taken, and 5 ml samples of culture were removed and harvested for reporter assays. Cells were grown for 30 hours at 70 ℃ for both the test of various P_slp_ truncations and the test of the chimeric P_slp_::RBS promoters. In all of the above experiments, each strain was grown in biological triplicate for every experimental condition.

### Measurement of *Cm*βgal reporter activity

To detect galactosidase activity in *A. bescii* cells, para-nitrophenol-β-D-galactopyranoside (pNPβGal), obtained from TCI chemicals, was used as colorimetric substrate. Substrate solutions contained 5 mM pNPβGal dissolved in 100 mM sodium acetate pH 5.5 buffer. β-galactosidase reporter assays were carried out as previously described in detail for *Cm*βgal^22^. Briefly, *A. bescii* cells were harvested and prepared in 100 mM pH 5.5 sodium acetate buffer before being heat-treated for 10 minutes at 90 ℃. Prepared cells were then added to the substrate solution, incubated for 10 minutes at 90 ℃, after which all reactions were immediately quenched with addition of 1M sodium carbonate. The absorbance at 405 nm (A405) of 100 µl of each reaction was measured in a flat-bottomed clear 96 well plate using a BioTek SynergyH1 microplate reader (Agilent). Normalized activity was also calculated as described previously including all controls to account for thermal background degradation of substrate and scattering from cellular debris, with normalization by the OD680 of the prepared *A. bescii* cells input into each reaction measured on a Nanodrop One C spectrophotometer blanked with 100 mM sodium acetate buffer^22^. All reaction conditions, including for each biological replicate, were performed in technical triplicate. Additionally, statistical significance of reporter activity driven by P_slp_ truncations relative to that of the native sequence was determined using a Brown-Forsythe and Welch ANOVA test on the collected data at the biological replicate level. Statistical significance was similarly determined between the reporter outputs of P_slp89_ and P_slp108_ by running a Welch’s T-test on these two data sets alone. Both statistical tests were performed in GraphPad Prism version 11.0 (GraphPad Software, Boston, Massachusetts USA).

### RNA extraction and 5’ RACE

RNA was isolated from *A. bescii* transformed with pJLG093, pJLG100, pJLG159, pJLG167, pJLG168, pJLG174, and pJLG215-19 grown on CM516K media for 16-18 hours. After growth, 30-40 ml cells were immediately pelleted at 6000 x g for 10 minutes, and frozen at -80 ℃ after removal of the supernatant. Cell lysis was carried out as previously described^13,22^ after which RNA was purified using the Monarch^®^ Spin RNA Isolation Kit (New England Biolabs) with the on-column DNase I treatment step. RNA concentrations were quantified using a Nanodrop One spectrophotometer (Thermo Scientific). 5’ RACE reactions were carried out with these RNA samples using the Template Switching RT Mix from New England Biolabs following the manufacturer’s recommended protocol for 5’ RACE. The reverse transcription and template switching step were carried out with the template switching oligonucleotide JLG_TSO1 (**Table S8**) purchased from Integrated DNA Technologies and the RT primer JLG243 (**Table S8**). Subsequent PCR amplification of cDNA used primers JLG242 & 244 (**Table S8**). After purification using the Clean and Concentrator-25 kit (Zymo Research), PCR products were quantified on a Nanodrop One spectrophotometer (Thermo Scientific), before being Sanger sequenced (Azenta Genewiz) with primer JLG218 (**Table S8** & **Supplementary File S1**). Alignment of this sequencing with the associated reporter construct and template switching oligonucleotide was used to determine putative transcriptional start sites (TSSs).

### Growth of AdhE expressing *A. bescii* and quantification of ethanol

*A. bescii* transformed with pSBS4, pJLG222, and pJLG224 were cultured in complex selective cellobiose media (CC516K). To begin the experiment, cultures were inoculated into anaerobic tubes as described above for the *Cm*βgal constructs. Cells were grown in biological triplicate for 48 hours at a lowered temperature of 65 ℃ to maximize ethanol titers^13^, after which samples were removed for analysis and measurement of the final OD680. Fermentation products (lactate, acetate, and ethanol) were quantified via HPLC. Samples were prepared for analysis by centrifugation at 21000 x g for 5 minutes, after which the supernatant was removed and passed through a 0.22 µm filter. Along with a filtered blank of CC516K media, these samples were analyzed using a 300 x 7.7 mm Agilent Hi-Plex H column heated to 50 ℃ and data was collected with an Agilent 1260 Infinity II Refractive Index detector also heated to 50 ℃. The mobile phase was 5 mM H_2_SO_4_ with a flow rate and injection volume of 0.6 mL/min and 10 µL respectively.

## Supporting information

Supporting Information .ab1 Sanger Sequencing

## Data Availability Statement

The data associated with this study are available in the article and its Supporting Information.

## Supporting Information

**Table S1**: reporter vectors, associated promoters, and promoter sequences

**Tables S2–S4**: reporter activity data for Figures 1b-d

**Table S5**: reporter activity data for Figure 2b

**Table S6**: reporter activity data for Figure 3c

**Table S7**: OD680 cell densities associated with Figure 4b

**Table S8**: oligonucleotide primer sequences

**Table S9:** DNA synthesis sequences

**Supplementary .ab1 Sanger sequencing files** from 5′ RACE supporting transcriptional start site assignments.

## Conflict of Interest

J.L.G. and J.M.C. are inventors on a provisional patent application related to genetic engineering of *A. bescii*. The remaining authors declare no competing financial interest.

## Author Contribution

**Joey L. Galindo:** Conceptualization, Methodology, Formal analysis, Investigation, Visualization, Writing - Original Draft

**Hansen Tjo:** Formal analysis, Investigation, Writing - Review & Editing

**Archit Srivastava:** Investigation, Writing - Review & Editing

**Miranda Harmon-Smith**: Project Administration

**Ian Blaby:** Conceptualization, Methodology, Supervision, Writing - Review & Editing **Jonathan M. Conway**: Conceptualization, Supervision, Resources, Writing - Review & Editing, Funding acquisition

## Funding

This work is supported by the A1531 Biorefining and Biomanufacturing program, project award no. 2026-67022-45882, from the U.S. Department of Agriculture’s National Institute of Food and Agriculture. This work was also supported by the Energy Research Fund administered by the Andlinger Center for Energy and the Environment at Princeton University and startup funds from the Department of Chemical and Biological Engineering at Princeton University.

## Acknowledgements

This project was enabled by DNA synthesis and plasmid construction conducted by the Joint Genome Institute (https://ror.org/04xm1d337) under proposal: https://doi.org/10.46936/10.25585/60012765. The U.S. Department of Energy Joint Genome Institute, a DOE Office of Science User Facility, is supported by the Office of Science of the U.S. Department of Energy operated under Contract No. DE-AC02-05CH11231. We would like to acknowledge Dr. Robert Evans at the Joint Genome Institute for carrying out the laboratory protocols associated with this synthesis project. Finally, we would also like to acknowledge Dr. Ryan Bing for his helpful advice regarding the growth characteristics and genetic manipulations of *A. bescii*, as well as the late Dr. Robert Kelly for his lifelong contributions to the study of extremophiles, especially *A. bescii*.

